# Determination of Bacterial Surface Charge Density Via Saturation of Adsorbed Ions

**DOI:** 10.1101/2020.09.29.318840

**Authors:** M.J. Wilhelm, M. Sharifian Gh., C.M. Chang, T. Wu, Y. Li, J. Ma, H.L. Dai

**Affiliations:** Temple University, United States

## Abstract

Bacterial surface charge is a critical characteristic of the cell’s interfacial physiology that influences how the cell interacts with the local environment. A direct, sensitive, and accurate experimental technique capable of quantifying bacterial surface charge is needed to better understand molecular adaptations in interfacial physiology in response to environmental changes. We introduce here the method of second harmonic light scattering (SHS) which is capable of detecting the number of molecular ions adsorbed as counter charges on the exterior bacterial surface, thereby providing a measure of the surface charge. In this first demonstration, we detect the small molecular cation, malachite green, electrostatically adsorbed on the surface of representative strains of Gram-positive and Gram-negative bacteria. Surprisingly, the SHS deduced molecular transport rates through the different cellular ultra-structures are revealed to be nearly identical. However, the adsorption saturation densities on the exterior surfaces of the two bacteria were shown to be characteristically distinct. The negative charge density of the lipopolysaccharide coated outer surface of Gram-negative *E. coli* (8.7±1.7 nm^−2^) was deduced to be seven times larger than that of the protein surface layer of Gram-positive *L. rhamnosus* (1.2±0.2 nm^−2^). The feasibility of SHS deduced bacterial surface charge density for Gram-type differentiation is presented.

**STATEMENT of SIGNIFICANCE:** Bacterial surface charge density is an important physiological characteristic which determines how the cell interacts with its local environment. Directly measuring the surface charge density, however, is experimentally non-trivial. In this work, we report an experimental method, second harmonic light scattering, that can directly and accurately quantify the surface charge density of individual living bacteria. This is achieved by measuring the number of molecular ions electrostatically adsorbed on the exterior cellular surface as counter charges. It is found that the negative charge density of a representative Gram-negative bacterium is 7 times larger than a representative Gram-positive bacterium. It is suggested that this disparity of surface charge density can be exploited as a basis for Gram-classification of bacteria.

## INTRODUCTION

The bacterial cell envelope defines the boundary physically separating the cell interior from the extracellular environment and plays a critical role in the physiology of the cell. Most significantly, the envelope acts as a mediator for exchange of vital nutrients and signaling molecules as well as for adhesion properties of the cell.(1) With rare exception, the surface of the bacterial outer envelope carries a net negative charge.(2) This stems primarily from the ionized phosphate and carboxyl groups localized on the bacterial surface.(2) The molecular composition of the exterior surface evolves over the course of normal development and may change in response to acute changes in environmental factors.(3) As the molecular composition changes, so does the surface charge density. For instance, physiologically induced variations of surface charge have been implicated in resistance mechanisms in response to antimicrobial peptides.(4) Bacterial surface charge and likewise the degree of hydrophobicity greatly influence how bacteria interact with their local environment. In addition to the uptake and release of molecules, electrostatic forces have been identified as critical initiation events for the formation of bacterial biofilms.(5) Overall, as interfacial physiology is vital to the continued well-being of bacteria, a significant fraction of their metabolic energy is devoted to maintaining the chemical composition of the exposed outer cellular surface.(1) Developing a better understanding of bacterial interfacial physiology therefore requires experimental methodology capable of accurately characterizing cellular surface charge density.

A number of different experimental approaches for determining surface charge have been developed. They include microelectrophoresis, electrostatic interaction chromatography, biphasic partitioning, isoelectric equilibrium analysis, and electrophoretic light scattering.(1) Most modern approaches typically rely upon characterization of the bacterial zeta potential (i.e., the electrical potential of the interfacial region between the cell surface and the local aqueous environment) to infer the surface charge.(1) Technically, the zeta potential is an observable which results from a convolution of the surface charges and associated solvation shells. The surface charge can only be theoretically deduced from this observable. Herein, we propose a direct measure of bacterial surface charge by quantitatively measuring the saturation concentration of electrostatically adsorbed counter ions using optical nonlinear light scattering.

We have previously demonstrated the use of time-resolved second-harmonic light scattering (SHS) for monitoring molecular interactions (e.g., surface adsorption and membrane transport) with living cells.(6) SHS is a nonlinear optical technique and is inherently surface sensitive. It is based upon the physical phenomenon, second-harmonic generation (SHG), whereby a portion of an incident light of frequency ω is converted to 2ω after interacting with a material.(7, 8) The physics of SHS and its application for probing the surface of colloidal particles have been comprehensively detailed in the literature.(9–11) Briefly, any molecule lacking center of inversion symmetry is SHG-active and is therefore capable of exhibiting a non-zero SHS response. However, a disordered ensemble of such molecules, for instance in a liquid solution, will not produce a coherent SHS response due to destructive interference from oppositely oriented nearest neighbor molecules. Nevertheless, if these molecules adsorb onto a surface or interface, they can align with one another and hence generate a coherent SHS response.

This characteristic mechanism has previously been employed to monitor molecular adsorption and transport across phospholipid membranes in biomimetic model liposomes(12–19) and living cells.(20–28) When SHG-active molecules adsorb onto the exterior surface of the membrane, they align with one another due to the similarity of the interaction driving the adsorption process. This oriented ensemble of SHG-active molecules gives rise to a coherent SHS response, the intensity of which scales as the square of the molecular surface density. After diffusing across the membrane, the molecules can then adsorb onto the interior leaflet of the membrane. This oppositely oriented ensemble of molecules (relative to the exterior leaflet) results in destructive interference of the time-dependent SHS signal. Consequently, the sequential rise and decay of SHS signal is quantitatively characteristic of the molecular adsorption and transport kinetics across a membrane. This interpretation has been validated in prior studies using optical brightfield transmission microscopy.(20, 29)

We now demonstrate that SHS can be used as a means of quantifying the surface charge density in ensembles of living cells. Specifically, as a representative example, we measure the surface charge of two very different strains of bacteria: one Gram-positive and one Gram-negative. As depicted in **Figure 1**, these two cell types are structurally and compositionally dissimilar. In general, the cell envelope of Gram-negative cells consists of a pair of phospholipid membranes, an exterior outer membrane (OM) and an inner cytoplasmic membrane (CM). These two membranes are separated by a thin diffusion barrier known as the peptidoglycan mesh (PM). Conversely, Gram-positive cells have only a single phospholipid membrane, the inner CM, which is surrounded by a substantially thicker PM. Additionally, Gram-positive cells can also possess an external shell of crystallized protein denoted as the surface, or S-layer.(30, 31)

**Figure 1.**
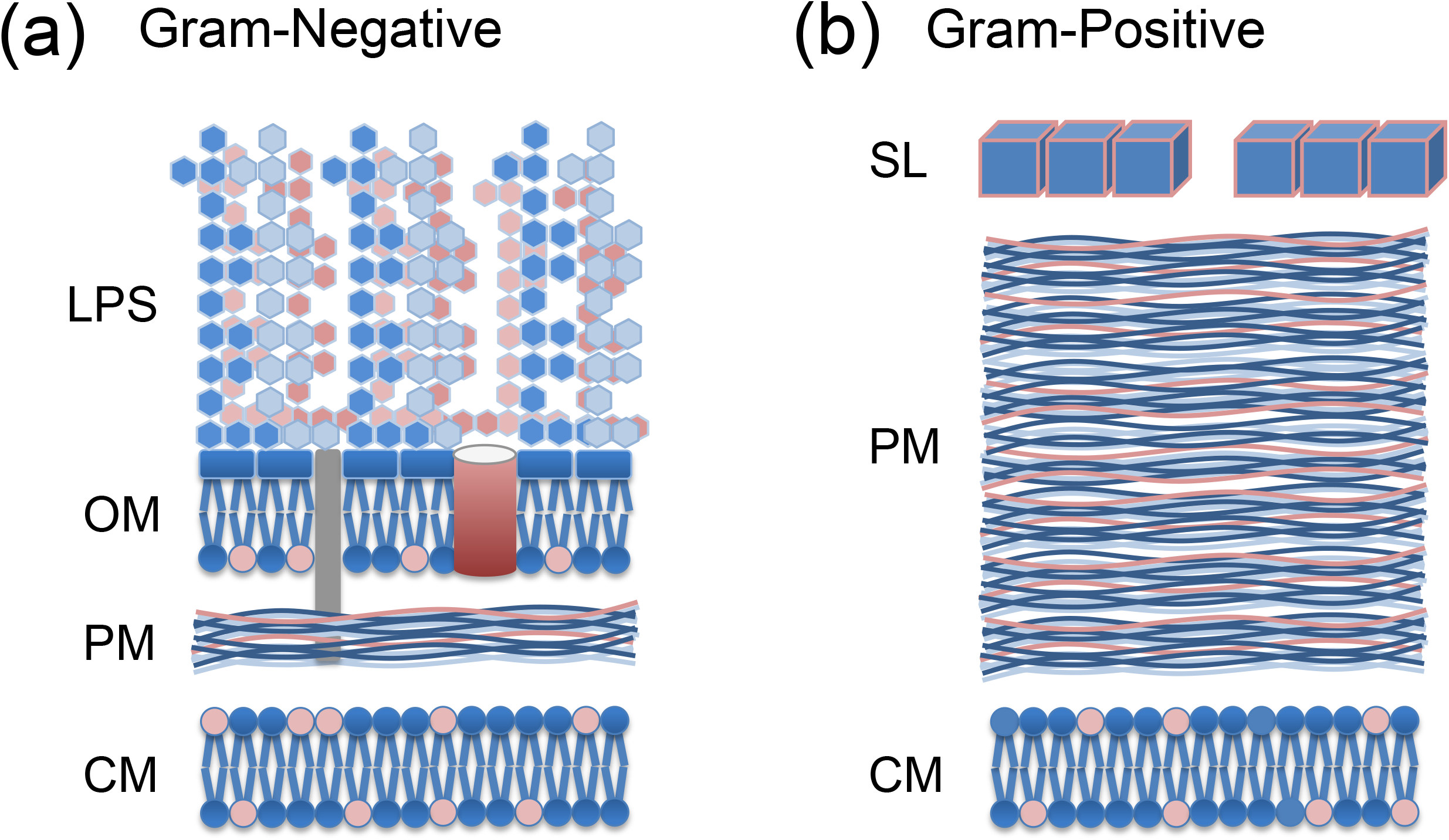
Comparison of the general ultra-structure of (a) Gram-negative and (b) Gram-positive bacteria.

Based upon these characteristic structural and compositional differences in cell envelopes (**Figure 1**), it is reasonable to speculate that the external surfaces of these two cell types should exhibit distinct molecular adsorption behaviors. Specifically, whereas Gram-negative cells are covered in long polyanionic lipopolysaccharide (LPS) hairs (and are therefore rough and porous), Gram-positive cells are coated with a comparatively smooth crystalline protein shell. While the acidic amino acids of the S-layer also yield a net negative surface charge,(32) the three dimensional nature of the LPS surface imparts a significantly higher surface charge density. We therefore postulate that these two general cell types should exhibit quantitatively differentiable characteristics based upon the achievable saturation density of adsorbed ions on their respective external surfaces. In particular, by quantifying both the ion saturation density and the available surface area of the cells, it is feasible to deduce the bacterial surface charge density.

In order to demonstrate proof-of-concept, we examined the interactions of the SHG-active molecular ion, malachite green, with two different strains of bacteria: Gram-positive *Lactobacillus, L. rhamnosus* (R0011) and Gram-negative *Escherichia, E. coli* (mc4100). Malachite green (MG, C_25_H_25_N_2_) is a small (650 Dalton) hydrophobic molecular dye that belongs to the family of triphenyl methanes. It has a pKa of 6.9 and under physiological conditions, a significant fraction of the MG population exists as cations.(33) The MG cation has an electronic absorption band near 400 nm and consequently its second order nonlinear polarizability is resonantly enhanced when exposed to fundamental light of 800 nm.

Adsorption of MG cation onto bacterial surfaces occurs primarily through an attractive electrostatic interaction. For each bacterial strain, we ran a series of SHS experiments in which we measured the time-dependent molecular uptake kinetics over a range of increasing MG concentrations. As demonstrated previously, the time-dependence allows selective isolation of the molecular interaction behavior (i.e., adsorption and transport) with the various bacterial interfaces.(6) For example, in the case of *E. coli*, MG first interacts with the exterior OM, then the PM, and finally the interior CM. The concentration dependent signal response at each interface enables us to construct Langmuir adsorption isotherms, which can then be used to quantitatively deduce the corresponding molecular saturation density. Finally, brightfield transmission microscopy and associated image analysis permits a quantitative measure of the surface area for each bacterial strain. Using the saturation density of adsorbed ions and the available surface area, we can deduce the surface charge density for each cell type examined.

## MATERIALS and METHODS

### A. Time-Resolved Second Harmonic Light Scattering

The specifics of our SHS experimental set-up has been detailed previously.(20, 34) Briefly, the output from a Ti:Sapphire laser (Coherent, Micra V, oscillator only, 800 nm, 150 fs pulse duration, 76 MHz repetition rate, 0.4 W average power, and 4 nJ pulse energy) was used as the fundamental light for SHG. The ultrafast laser pulses were used for the high peak intensity in order to achieve higher nonlinear optical signal, but with low pulse energy in order to minimize absorption induced damage to the sample. To minimize background noise from light scattering from interfaces, the fundamental light pulses interacted with the sample in a continuously flowing liquid jet, roughly 1-2 mm in diameter. The liquid sample was pumped from a 10 mL sample reservoir that was continuously stirred using a magnetic stirrer. The laser pulse was focused into the liquid jet with a waist diameter of roughly 40 μm and a Rayleigh length of 1.6 mm, yielding a focal volume of approximately 6 nL. The scattered signal was collected in the forward propagation direction defined by the fundamental light beam. The laser output has a >99% linear polarization along the horizontal direction. All polarizations of second harmonic light scattered after the sample was collected. A long-pass filter (Schott, RG695, >650 nm) was placed in front of the sample to block any higher harmonic signal produced by the preceding optics. Likewise, a band-pass filter (Schott, BG39, 340-610 nm) and a monochromator (1 mm entrance and exit slits, 400 ± 1 nm bandwidth) were used to isolate the second harmonic light at the detector. The SHS signal was detected with a photomultiplier (Hamamatsu, R585), pre-amplified (Stanford Research Systems SR4400), and processed using a correlated photon counting system (Stanford Research Systems, SRS SR400).

### B. Time-Resolved Brightfield Transmission Microscopy

Brightfield transmission microscopy was used to quantify the size and shape of individual bacteria. All images were acquired on a Leica DMRXE microscope using a 100× PlanApo objective lens that was coupled to a digital image capture system (Tucsen, model TC-3) and software controlled using TSView (OnFocus Laboratories, ver. 7). The field of view (FOV) covered an area of 60×50 μm^2^. The images were saved using the tagged image file format (TIFF). Cells were immobilized by pretreating the slides with poly-L-lysine. Image analysis was performed in ImageJ (National Institutes of Health, 1.43u).

### C. Sample Preparation

Colonies of *L. rhamnosus* (R0011) and *E. coli* (mc4100) were grown on Luria Broth agar (Sigma-Aldrich) plates. Bacterial samples were cultured (37°C with 150 RPM shaking) in Terrific Broth (Sigma Aldrich) to late-log/early-stationary phase, lightly pelletized by centrifugation, and washed three times in 1xPBS to remove waste and residual growth medium. Washed pellets were then re-suspended in 1xPBS (Sigma) to achieve working sample OD_600_ values of 0.5 (ca., 5.0×10^8^ cfu/ml). Concentrated stock solutions of MG were prepared by dissolution of the oxalate salt and used as obtained from the supplier (Sigma-Aldrich). The pH of the sample (MG and cells) was stable at 7.3.

## RESULTS & ANALYSIS

### A. Time-Resolved SHS Signal

**Figure 2** depicts representative normalized SHS signals corresponding to the interaction of MG (10 μM final concentration) with either Gram-positive (*L. rhamnosus*, violet circles) or Gram-negative (*E. coli*, pink circles) bacteria. Note that time has been plotted on a logarithmic scale in order to simultaneously highlight both the fast and slow molecular transport kinetics. Qualitatively, both traces appear very similar and can be assigned to a fast transport event followed by a second slower transport event. In general, the SHS signal rises due to increasing adsorption of MG onto a surface of the bacteria, while the subsequent signal decrease corresponds to the build-up of MG density on the opposite surface following transport across the cellular interface.(6) For *L. rhamnosus*, the fast event can be assigned as rapid transport across the numerous pores (2-8 nm diameter) which perforate the bacterial S-layer. This is followed by a second slower transport event as MG adsorbs onto and diffuses across the bacterial CM. For *E. coli*, the fast transport event is assignable to rapid transport across the outer membrane protein (Omp) channels in the OM. Likewise, this is followed by much slower direct diffusion of MG across the CM.

**Figure 2.**
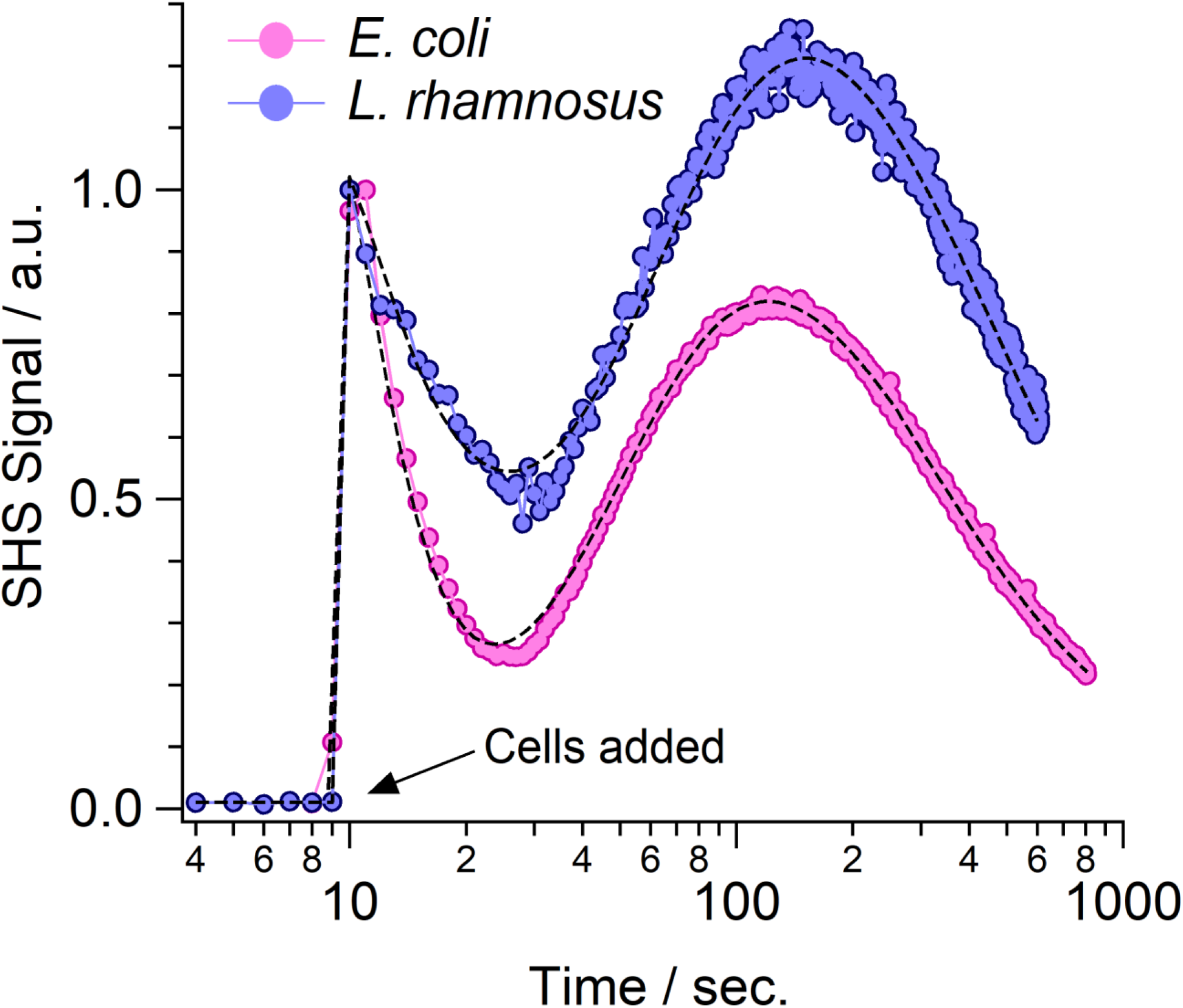
Representative time-resolved SHS signal following addition of bacteria (pink circles, *E. coli* and violet circles, *L. rhamnosus*) to solutions containing 10 μM MG. Bacteria were added to the system around t = 9 s. Dashed lines are best fit results based upon a kinetic model of molecular uptake.

### B. Molecular Transport Rates

The measured time-resolved SHS kinetic traces can be fit, using our previously developed kinetic model of molecular adsorption and transport, to determine the molecular transport rates for the various bacterial interfaces (i.e., outer membrane, peptidoglycan mesh, S-layer, and cytoplasmic membrane).(20–28) It is found that MG cation transport across the Omps in the OM and the pores in the S-Layer are similarly rapid and were deduced as 0.04 s^−1^ and 0.02 s^−1^, respectively. This is reasonable as transport across these interfaces involves an MG cation (whose longest dimension is roughly 1 nm) traversing comparatively massive holes of 2-8 nm in diameter. Likewise, direct transport across the hydrophobic core of the CM was found to be virtually identical for both bacterial strains, with a deduced rate of 2.0×10^−4^ s^−1^. Of greater interest are the transport rates across the PM, which were determined as 0.07 s^−1^ (*E. coli*) and 0.04 s^−1^ (*L. rhamnosus*). Given that Gram-positive bacteria have substantially thicker PM’s than Gram-negative bacteria, it is surprising that these rates are so similar and suggests that MG does not interact significantly with the PM.

### C. Langmuir Adsorption Isotherm for the Bacterial Outer Surfaces

**Figure 3** shows representative SHS signal traces following the addition of increasing concentrations of MG to a solution containing either *E. coli* (left) or *L. rhamnosus* (right). Here we focus only on the initial portion of the SHS response which corresponds to adsorption and rapid transport across the external interface of the bacteria (i.e., the OM for *E. coli* and the S-layer for *L. rhamnosus*). The MG concentration range was chosen in order to achieve saturation on the exterior surface of the bacteria. Coherent SHS signal is only produced from MG cations adsorbed on the bacterial surface. As the concentration of MG is increased, the resulting SHS signal likewise increases and scales as the square of the molecular surface density. However, once saturation is achieved (i.e., no further MG cations can adsorb on the bacterial surface), there is no further increase of the measured SHS signal. It is of interest to note that the saturation densities are clearly different for the two bacteria. Additionally, under conditions of saturated adsorption at the surface, roughly four times more SHS signal is measured for MG cations interacting with *L. rhamnosus* (ca. 80,000 counts/s) than *E. coli* (ca. 20,000 counts/s).

**Figure 3.**
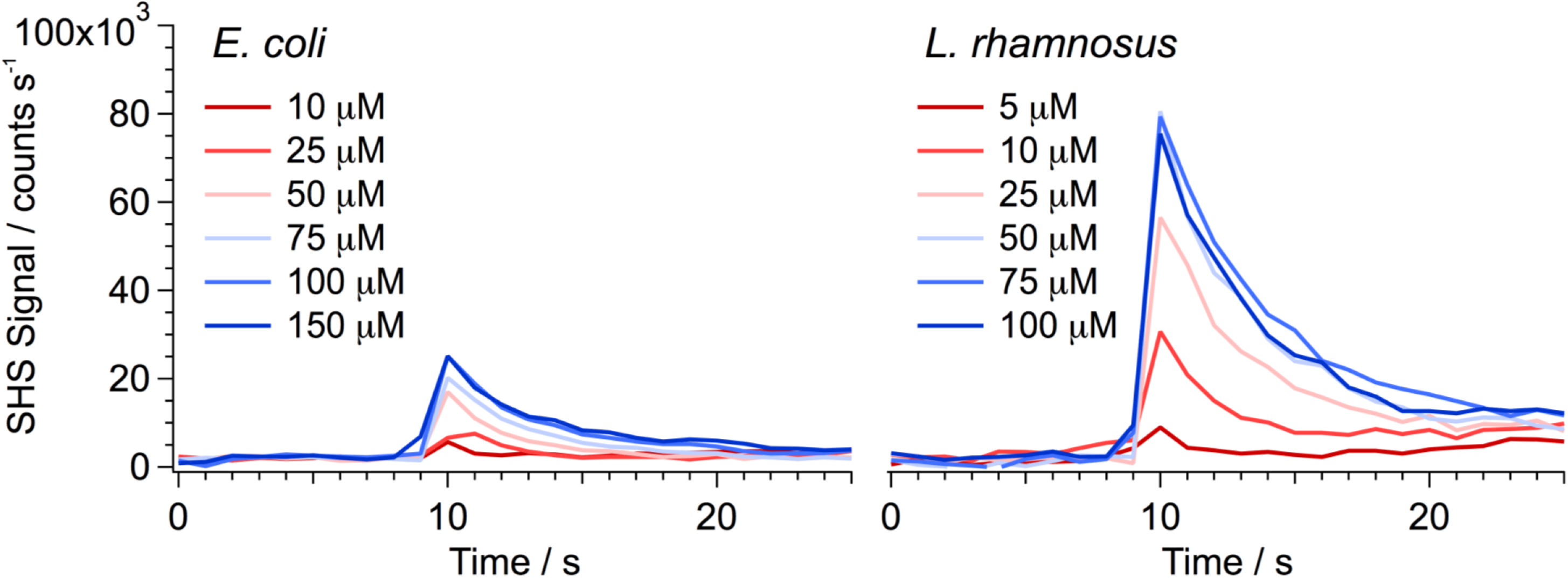
Representative time-resolved SHS signal following addition of bacteria (left, *E. coli* and right, *L. rhamnosus*) to solutions containing increasing concentrations of MG. Bacteria were added to the system around t = 9 s.

To determine the saturation density of MG cations adsorbed on the external surface of the bacteria, we measured Langmuir adsorption isotherms by plotting the SHS peak intensities (deduced from **Figure 3**) as a function of the applied MG concentration. Note that the SHS intensity is proportional to the square of the MG cation density at the surface. **Figure 4** shows a comparison of the Langmuir adsorption isotherms for the OM of *E. coli* (pink circles) and the S-layer of *L. rhamnosus* (violet circles).

**Figure 4.**
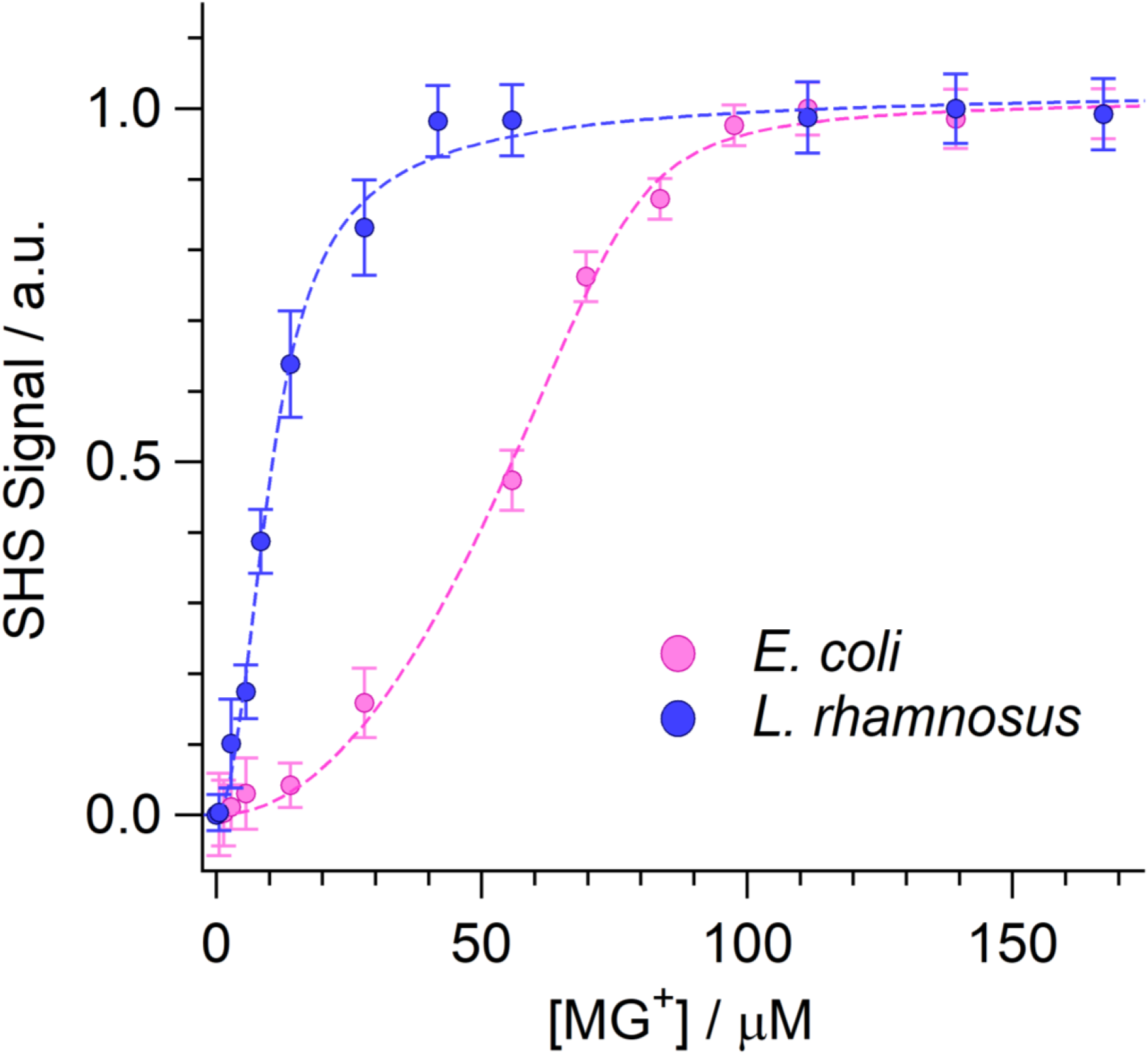
Langmuir adsorption isotherms for MG interacting with the exterior surfaces of *L. rhamnosus* (violet circles) and *E. coli* (pink circles). Dashed lines represent best fit results to the modified Langmuir adsorption model.

In order to properly analyze the Langmuir isotherm, we must take into account the fact that in our experiments, only cationic MG yields a resonantly enhanced SHS response and therefore produces the measured signal.(8) At pH = 7.3, MG is only about 30% cationic. As the MG cation concentration increases, the magnitude of the measured SHS signal likewise increases. Once the surface of the bacteria becomes fully saturated with cations, the resulting signal levels out to a plateau. A nonlinear least squares fit of the Langmuir isotherm using the modified Langmuir adsorption model(35) (i.e., which includes the constraint that the total quantity of MG in the solution and on the bacterial surfaces is constant) permits determination of the saturation surface density (N_max_) as well as the equilibrium adsorption constant, from which the free-energy of adsorption (ΔG_ads_) can be deduced.(36)

### D. Adsorption Free Energy and Saturation Density of MG Cations on the Bacterial Outer Surfaces

The magnitude of the deduced ΔG_ads_ for *E. coli* (−10.7±0.3 kcal/mol) is slightly larger compared to *L. rhamnosus* (−10.2±0.3 kcal/mol), but both values are consistent with an electrostatically driven adsorption process.(21) Despite the similarity of the adsorption free energies, the concentrations at which surface saturation occurs for the exterior surface for the two bacteria are an order of magnitude different. For *L. rhamnosus,* saturation occurs at a modest MG cation concentration of 9.6 μM, while *E. coli* saturates at a significantly higher concentration of 77.2 μM. This suggests that there is eight times more MG on the exterior surface of *E. coli* compared to *L. rhamnosus*. With the cation saturation concentrations determined, we next quantified the sizes of the two strains of bacteria so that surface charge density could be calculated.

In order to determine the sizes of these two strains of bacteria, we imaged the cells using brightfield transmission microscopy (**Figure 5**). Specifically, the available surface area (S_A_) of each cell type was estimated by fitting the shape of the bacteria to an ellipse for the determination of their length (major axis) and width (minor axis). This process was repeated for more than 325 cells of each strain. The distributions of the measured major and minor axes for each strain are plotted in **Figure 5**. As revealed in the images (and quantitatively in the plots), these strains of *E. coli* and *L. rhamnosus* are very similar in size, roughly 2.8 μm long by 1 μm wide. In order to determine the average surface area of the cells, we modeled the bacteria shape as a cylinder between two halves of a sphere (**Figure 5C**). Within this framework, the cellular surface area can be estimated. It is found that *E. coli* has a slightly larger average surface area of 11.0±1.5 μm^2^ compared to 10.1±1.4 μm^2^ for *L. rhamnosus*. Using the saturation concentrations determined in the Langmuir adsorption isotherms (**Figure 4**), the known cell densities of the samples (5×10^8^ cells/ml), and the surface areas determined in the image analysis (**Figure 5**), the saturation densities for the two bacterial strains can be deduced as: 8.7±1.7 nm^−2^ (*E. coli*) and 1.2±0.2 nm^−2^ (*L. rhamnosus*).

**Figure 5.**
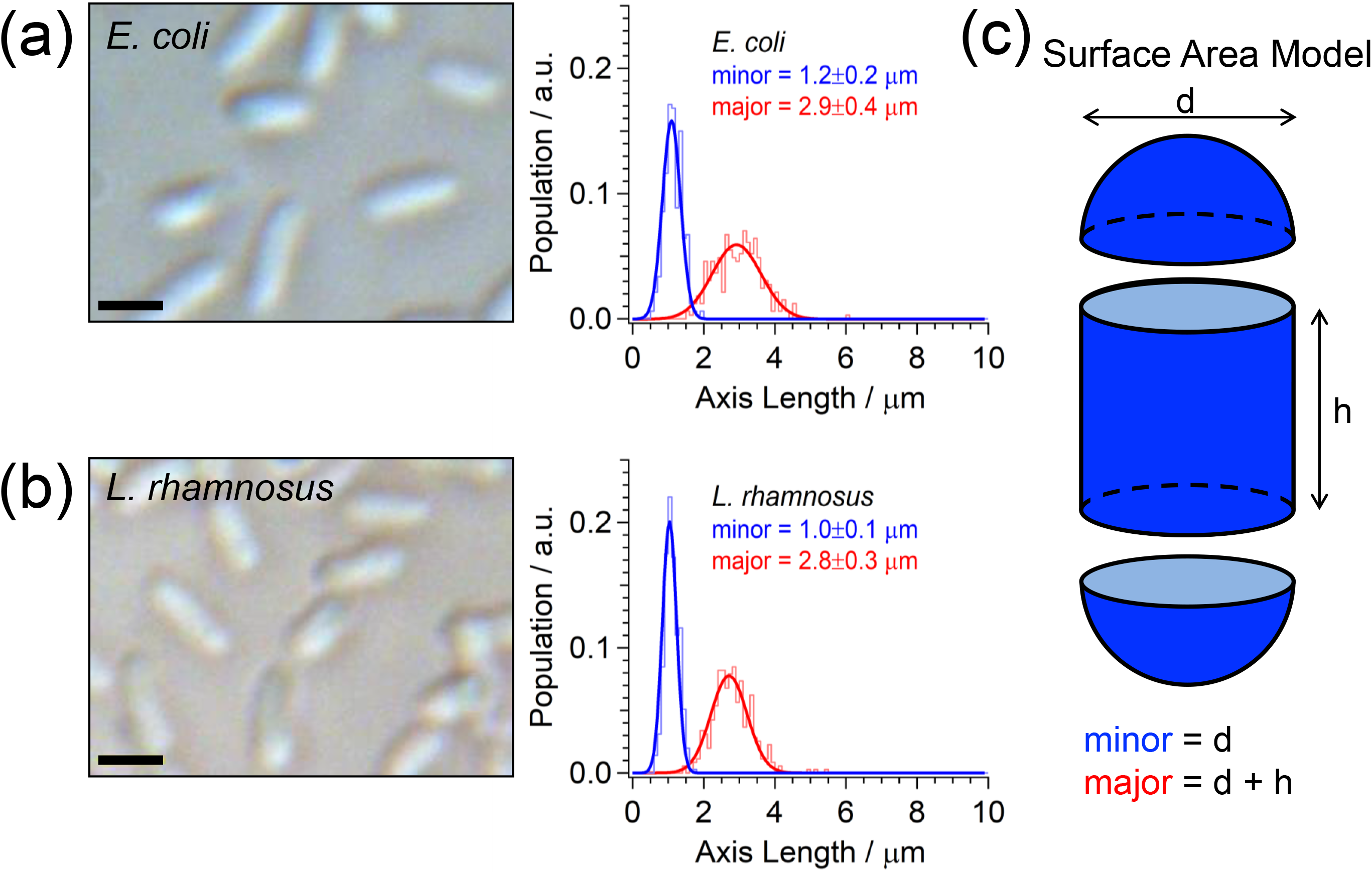
Brightfield transmission images and measured size distributions of the major (red) and minor (blue) length axes of (a) *E. coli* and (b) *L. rhamnosus*. Scale bars depict 2 μm. (c) Surface area (S_A_) modeled as the combined exposed surface of a sphere and a cylinder.

## DISCUSSION

Over a series of prior studies, we have extensively quantified the molecular uptake kinetics of the molecular MG cation in the proto-typical Gram-negative bacteria, *E. coli*.(6) Using the surface sensitive time-resolved SHS technique, we have experimentally deduced the adsorption and transport kinetics for MG cations interacting with the bacterial OM, PM, and CM. As an extension of this work, we now examine the various molecular interactions of MG with a representative Gram-positive bacterium, *L. rhamnosus*. When considered together, this body of work helps to elucidate how the known structural differences (which distinguish Gram-positive and Gram-negative cells) influence molecular adsorption and transport. In the same vein, the different molecular interactions could provide an alternative means for differentiating these two cell types.

### A. Invariance of Molecular Transport through Different Ultra-structures

Gram-positive and Gram-negative cells have very different cellular ultra-structures. It is expected that these different cell types should exhibit distinguishable molecular uptake kinetics as the molecule necessarily interacts with different local environments along the way. However, as revealed in **Figure 2**, at least for the MG cation, this is decidedly not the case. Despite the numerous structural differences, the transport kinetics of MG were observed to be nearly invariant for these two cell types. A possible explanation for this behavior is that while the chemical composition of the cell wall components is distinct for Gram-positive vs. Gram-negative cells, their general structures are topologically similar (**Figure 1**). For instance, as shown in **Figure 2**, both cell types exhibit an initial rapid transport event through the outermost cellular interface (i.e., the S-layer for Gram-positive and the OM for Gram-negative). While compositionally different, they are both hydrophobic barriers perforated with large (i.e., 2-8 nm wide) water filled channels. From the perspective of a 1 nm wide reporter molecule, both of these interfaces are therefore quite similar. Likewise, beyond the outermost cellular interface, both cell types have a PM: a thick PM in Gram-positive cells and a substantially thinner PM in Gram-negative cells. Similar to the S-layer, the PM is known to be regularly perforated with comparatively massive pores, 7-12 nm wide.(37) As MG is so much smaller than the PM pores, it appears to be able to rapidly cross the PM regardless of thickness.

Likewise, both Gram-positive and Gram-negative cells have an interior CM which hinders the passive uptake of purely hydrophilic compounds. As demonstrated previously in numerous liposome based studies,(12–19) the MG cation is readily able to passively diffuse across phospholipid membranes. It is therefore reasonable that the MG cation should likewise cross the bacterial CM’s of both Gram-positive and Gram-negative cells and at a comparable rate.

### B. MG Cation Adsorption Density on Bacterial Surfaces

While molecular transport kinetics of MG seems altogether indifferent to the general ultrastructure of Gram-positive vs Gram-negative cells, its surface adsorption density is vastly different for these two cell types. The Gram-negative cells have a maximum surface adsorption density which is seven times larger compared to the Gram-positive cells.

How can we reconcile the vastly different adsorption densities for these two distinct strains of bacteria? For *L. rhamnosus*, a maximum adsorption density of 1.2±0.2 nm^−2^ actually seems quite reasonable. MG is roughly 1 nm wide and could therefore conceivably sit within a 1×1 nm^2^ area. The exterior surface of the bacterial S-layer (Gram-positive) is remarkably different compared to the LPS coated OM (Gram-negative). The bacterial S-layer is a homogeneous protein wall, self-assembled into a crystalline lattice with regularly spaced pores of distinct symmetry (e.g., oblique, square, hexagonal).(31) Compared to the rough 3D surface structure of the LPS, the S-layer is comparatively smooth and reasonably described as a 2D flat surface.(32) Furthermore, it is known that the S-layer is composed predominantly of acidic amino acids, which impose a net negative charge on the resulting S-layer.(32) Given these constraints, a maximum achievable adsorption density of a single MG cation per nm^2^ is quiet reasonable. Further, based on the deduced adsorption free energy, which is similar to those determined for adsorption driven by an attractive electrostatic interaction, we can conclude that the negative charge density on the *L. rhamnosus* outer surface is roughly one per nm^2^.

Conversely, the deduced *E. coli* adsorption density of 8.7±1.7 nm^−2^ seems high, at least in the context of a 2D planar surface. However, it is crucial to remember that, unlike Gram-positive cells, the exterior surface of Gram-negative bacteria is covered with long LPS hairs (**Figure 1**). It has previously been shown that the LPS inner core covers a rectangular area of roughly 0.8 nm^2^.(38) Given our deduced MG cation saturation density of 8.7±1.7 nm^−2^, this suggests an average net anionic charge of q = 7.3±1.8 for each of the LPS in our strain of *E. coli*. This value is reasonably consistent with previously deduced anionic charges for purified LPS isolated from strains of *S. minnesota* (q≥5), *E. coli* (q≥5), and *K. pneumoniae* (q=7).(39) Indeed, this is physically reasonable given the presence of numerous anionic phosphates localized on the inner core and Lipid A of the LPS.(38–40) Consequently, *E. coli* (and Gram-negative bacteria in general) should exhibit an overall higher cationic saturation density compared to Gram-positive cells due to the availability of this favorable electrostatic interaction with the poly-anionic LPS.

Understanding our experimental observations would not be complete without addressing the issue of the relative magnitudes of the SHS signal measured for both of the cell strains examined here. As revealed in **Figure 4**, MG surface saturation on *L. rhamnosus* yields roughly four times more signal compared to *E. coli*. This ratio seems to contradict the observation that there is seven times more MG per unit area on *E. coli* compared to *L. rhamnosus*. A plausible explanation for this behavior is the relative orientation of the MG ensemble on each of the two cell surfaces. Specifically, as depicted in **Figure 6**, it is likely that MG is more aligned with the surface normal on *L. rhamnosus*, as the molecular ion is expected to adsorb onto the negative charge sites of a comparatively flat surface. This adsorption geometry is expected to yield a larger coherent nonlinear polarization from the adsorbed ensemble of MG cations through constructive interference. In contrast, on the outer surface of *E. coli*, the MG ions adsorbed at the negative charge sites of the LPS are presumably more tilted toward the surface plane. The varying orientation within the bacterial surface plane may result in partial cancellation of the nonlinear polarizability and hence a smaller SHS signal.

**Figure 6.**
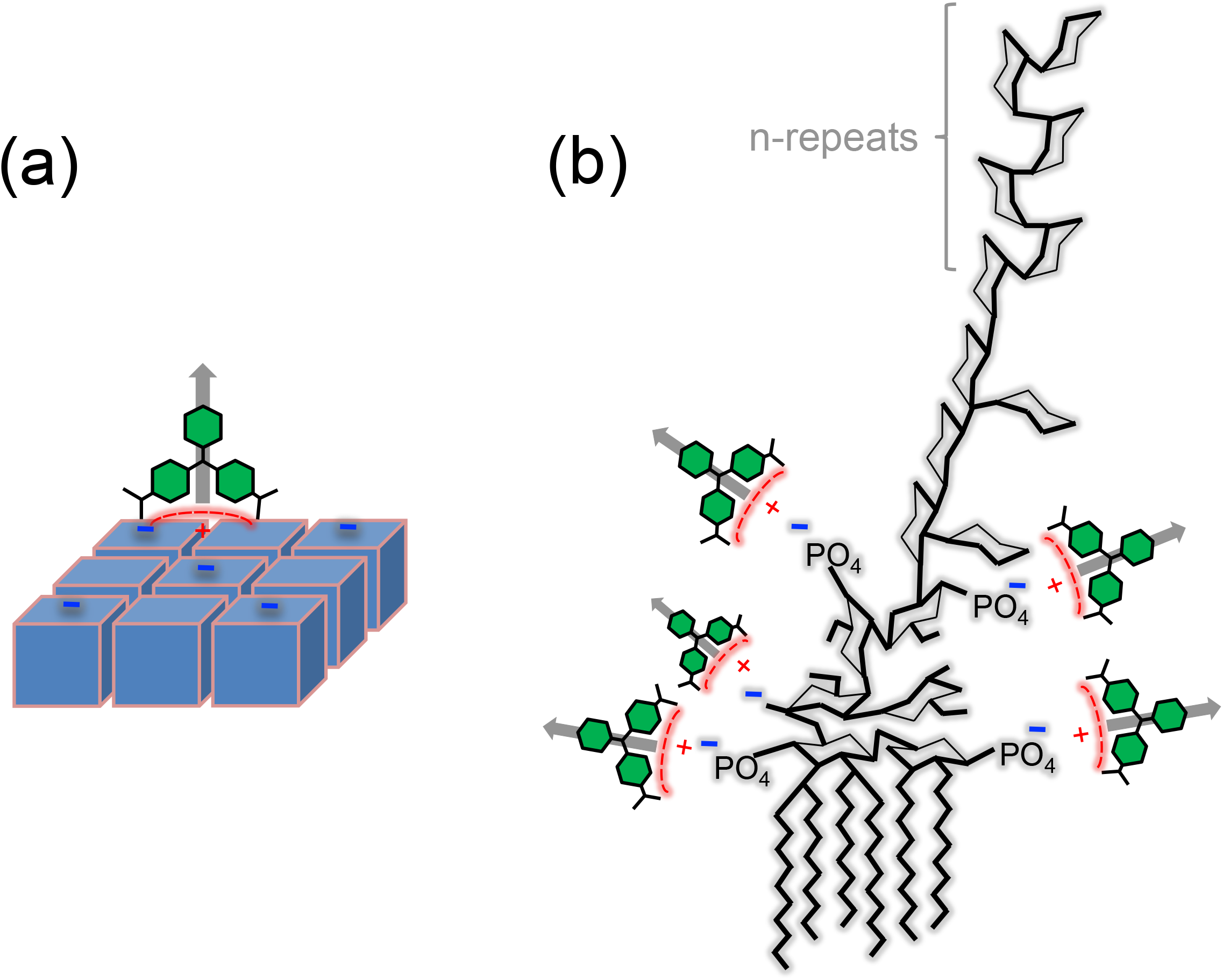
Schematic of the orientation of cationic MG adsorption on (a) the S-layer of a Gram-positive bacteria and (b) the LPS of a Gram-negative bacteria. Grey arrows indicate the molecular frame orientation of the hyperpolarizability of the MG cation.

### C. MG Cationic Adsorption Density for Gram Differentiation

The significant difference in the cationic MG adsorption saturation density between the two strains of bacteria can likely be exploited as an experimental basis for Gram-stain differentiation. We propose to simply measure the SHS-based Langmuir adsorption isotherms to establish Gram classification. Specifically, as shown in **Figure 4**, it is clear that the Gram-negative species requires a significantly higher concentration of the cation to fully saturate the exterior surface. It must be stressed, however, that these isotherms will vary as a function of cell density as well as the available surface area of the cell. For comparative analysis, it is therefore crucial to ensure that a constant sample OD is used. Here, we used a cell density of ca. 10^8^ cells/ml which was sufficiently high to yield a strong SHS signal, but still low enough to exclude effects of cell-cell interactions. Nevertheless, in the interest of greater certainty, the charge density of the cells should be determined. For Gram-positive cells, this corresponds to values of roughly 1 cation per nm^2^. Conversely, for the polycationic LPS of Gram-negative cells, this yields much larger values of >5 cations per nm^2^.

## CONCLUSIONS

We have applied time-resolved second-harmonic light scattering to characterize the uptake of a molecular cation, malachite green, in representative strains of Gram-positive and Gram-negative bacteria. Despite distinct compositions and cellular ultra-structures, the characteristic molecular transport kinetics for these two cell types were observed to be remarkably similar. It was argued that this stemmed primarily from the similarity of the general topology of the two ultra-structures.

The second-harmonic light scattering observations also allowed the measurement of Langmuir adsorption isotherms of MG cations on the outer surfaces of the two bacteria. Examination of the saturation adsorption density on the exterior surface of the two cell types revealed strikingly different behavior. Specifically, the comparatively smooth protein S-layer of the Gram-positive cell exhibits a relatively small adsorption density (ca. 1.2±0.2 nm^−2^), whereas the rougher LPS covered surface of the Gram-negative cells have a significantly larger adsorption density (ca. 8.7±1.7 nm^−2^). As the adsorption free energies determined from analyzing the Langmuir isotherms indicate that the adsorption is driven by attractive charge-charge interaction, the measure adsorption densities correspond to the negative charge densities at the two respective outer surfaces. It was suggested that this characteristic difference in saturation adsorption densities could be employed as an experimental metric for Gram classification.

## AUTHOR CONTRIBUTIONS

MJW and HLD designed the study. MSG conducted the experiments. MJW, CMC, TW, and YL analyzed the data. MJW, JM, and HLD interpreted the results. MJW and HLD wrote the manuscript.

## ACKNOWLEDGEMENTS

This work was supported in part by the Air Force Office for Scientific Research, under Grant Number FA9550-15-1-0213.

